# Cyanobacterial sigma factor controls biofilm-promoting genes through intra- and intercellular pathways

**DOI:** 10.1101/2023.12.14.570974

**Authors:** Shiran Suban, Sapir Yemini, Anna Shor, Hiba Waldman Ben-Asher, Orly Yaron, Sarit Lampert, Eleonora Sendersky, Susan S Golden, Rakefet Schwarz

## Abstract

Cyanobacteria frequently constitute integral components of microbial communities known as phototrophic biofilms. These assemblages are not only widespread in various environmental contexts but also hold significant industrial relevance. Nevertheless, the governing elements responsible for cyanobacterial biofilm development have remained elusive. This study, which employs the model cyanobacterium *Synechococcus elongatus* PCC 7942, demonstrates that the RNA polymerase sigma factor SigF1, but not its paralog SigF2, is required for a biofilm-suppression mechanism that operates in this organism. Comprehensive transcriptome analyses identified distinct regulons under the control of each of these sigma factors. Additional data indicate that SigF1 regulates biofilm through its involvement in transcriptional induction of genes that include those for the primary pilus subunit: *sigF1* inactivation both prevents pilus assembly and abrogates secretion of a biofilm inhibitor. Consequently, expression is significantly upregulated for the *ebfG*-operon that encodes matrix components and the genes that encode their corresponding secretion system. Thus, this study uncovers a basic regulatory component of cyanobacterial communal behavior. Elevated expression of biofilm-promoting genes in a *sigF1* mutant supports an additional layer of regulation by SigF1 that operates via an intracellular mechanism.

## Introduction

Sigma factors play a crucial role in bacterial regulation by conferring the specificity of the RNA polymerase core complex for specific promoter sequences, facilitating transcription initiation. In heterotrophic bacteria these proteins are classified into two evolutionary and structurally unrelated families: the σ54 and the σ70 families; however, no σ54 homologs have been identified in cyanobacteria [1]. The σ70 family comprises group 1, so-called housekeeping sigma factors, as well as alternative sigma factors that dictate transcription of whole sets of genes, referred to as regulons [2, 3]. Control of an entire regulon by a single sigma factor facilitates immediate response to environmental cues [4–7].

Cyanobacterial alternative sigma factors are associated with a large variety of cellular responses [1, 8–12]. For example, sigma factors of the filamentous cyanobacterium *Nostoc punctiforme* promote development of motile filaments known as hormogonia [13]. In the filamentous cyanobacterium *Anabaena* PCC 7120, genes *sigC*, *sigE*, and *sigG* are upregulated in heterocysts, specialized cells involved in N_2_ fixation [14]. Additionally, SigF, SigG, and SigH, which belong to group 3-type σ70 factors, are required for the low-temperature growth of the unicellular marine cyanobacterium *Synechococcus* PCC 7002 [15]. In *Synechocystis* PCC 6803, mutants in *sigF* are impaired in inducing salt-stress proteins [16] and pilus assembly [17–19]. Furthermore, a *sigF*-deletion mutant of this cyanobacterium exhibits impaired vesiculation capacity compared to the wild-type [20].

Recent genetic screening implied involvement of a SigF-homolog of the freshwater unicellular cyanobacterium *Synechococcus elongatus* PCC 7942 in regulation of biofilm formation [21]. Previous studies from our group have identified a biofilm-suppression mechanism in this cyanobacterium, which relies on extracellular inhibitor(s) that repress transcription of the *ebfG*-operon. This operon encodes four proteins that enable biofilm formation (Fig. 1; also see [22–24]). A recent study revealed cell specialization in expression of this operon [25], the products of which are secreted by a type-I like secretion system [24, 26]. Additionally, EbfG1-3 are prone to form protein fibers, while EbfG4 acts as both a cell surface and a matrix protein [25]. These findings indicate that a subpopulation of cells provides “public goods” to support biofilm development by the majority of the cells.

**Figure 1:**
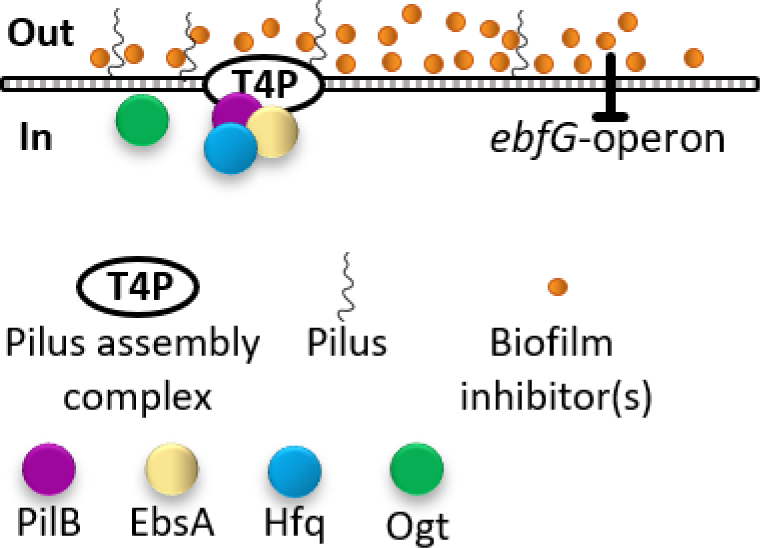
Expression of genes that enable biofilm formation is governed by extracellular inhibitor(s). PilB – assembly ATPase of the type IV pilus (T4P) assembly complex. EbsA – essential for biofilm suppression protein A. Hfq – homolog of RNA chaperone. Ogt – glycosyltransferase that glycosylates the pilus subunit PilA.

Cumulative evidence indicates that impairment of the type IV pilus assembly complex (T4P) disrupts the suppression mechanism leading to biofilm development (Fig. 1; also see [21, 22, 27, 28]), and affects the exoproteome, consistent with a shared complex serving in type IV pilus formation and type II secretion (cite Yegorov) Intriguingly, the RNA chaperone Hfq and a conserved cyanobacterial protein called EbsA (<u>e</u>ssential for <u>b</u>iofilm <u>s</u>uppression protein A) are part of the cyanobacterial T4P complex [28, 29]. The glycosyl transferase Ogt, responsible for modifying the pilus subunit PilA, is required for pilus formation and biofilm-suppression [30]. However, the components that regulate the expression of genes related to biofilm development have not yet been identified. This study assigns a pivotal role for a sigma F homolog of *S. elongatus* in the biofilm suppression process. Furthermore, the data demonstrate the involvement of this SigF in the regulation of biofilm-promoting genes through both inter-and intracellular pathways.

## Results and Discussion

### Inactivation of sigF1 results in biofilm development

A recent study showed that the inactivation of the gene Synpcc7942_1510, which encodes a homolog of the Sigma F factor known as SigF1, results in biofilm development [21]. Briefly, a pooled barcoded transposon library, RB-TnSeq, was grown, biofilms were formed, and sequencing of the barcodes indicated the abundance of each mutant in both the biofilm and the planktonic fractions. Data analysis revealed that Synpcc7942_1510 mutants are significantly over-represented in the biofilm, indicating the potential involvement of this gene in biofilm suppression. To characterize the specific role of SigF1 in biofilm suppression, the gene was inactivated by transformation with an allele disrupted by the *Mu* transposon (Fig. 2A and Table S1). The resulting mutant, SigF1::Mu, was then grown separately to observe its biofilm-forming ability.

**Figure 2:**
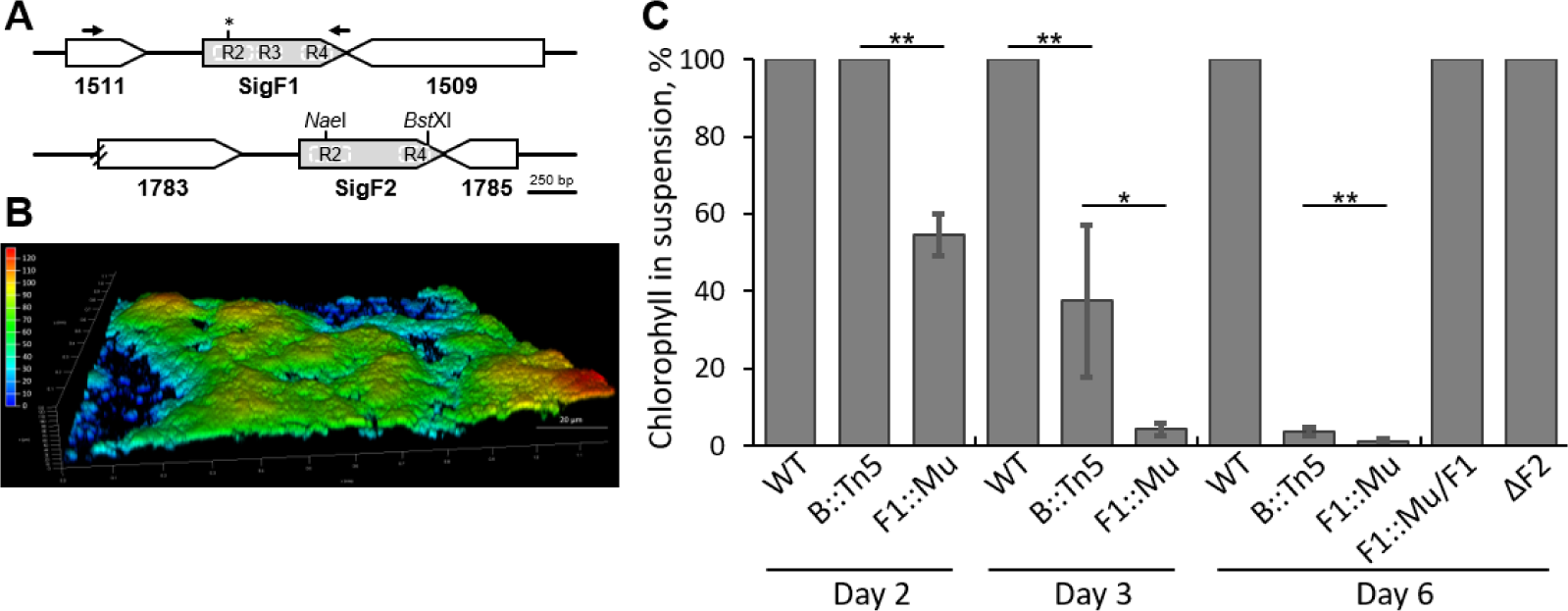
SigF1 is essential for the biofilm self-suppression mechanism. **A.** Genomic region of genes encoding homologs of sigma F factors of *S. elongatus*: synpcc7942_1510 and synpcc7942_1784. R2, R3 and R4 indicate particular domains of the SigF proteins. The asterisk denotes the insertion site of the Mu transposon in *sigF1*. NaeI and BstXI sites were used to construct the deletion mutant of *sigF2*. Arrows denote the primers used to PCR-amplify a DNA fragment for complementation. **B.** Image of SigF1::Mu biofilm using confocal fluorescence microscopy. Imaging is based on autofluorescence (excitation at 630 nm and emission at 641 to 657 nm). The color scale represents biofilm depth. **C.** Assessment of biofilm development by measurement of the percentage of chlorophyll in suspended cells. Robust biofilm development is manifested in low percentage of chlorophyll in suspended cells. Strains analysed: WT; the biofilm-forming strains in which *pilB* and *sigF1* were inactivated (B::Tn5 and F1::Mu, respectively); F1::Mu complemented with SigF1 (F1::Mu/F1) and a deletion mutant of *sigF2* (ΔF2). Data represent averages and standard deviations from 3 independent biological repetitions (with 3 technical repeats in each). Asterisks denote significance (t-test, two tails, two-sample assuming unequal variances. * p<0.001; ** p<5E^-5^).

Confocal fluorescence microscopy demonstrated the presence of robust biofilms of SigF1::Mu (Fig. 2B). Additionally, quantification of biofilm formation was achieved by measuring the relative amount of chlorophyll in the planktonic fraction of the culture[31]. On the second day of growth, SigF::Mu initiated biofilm formation, with approximately 60% of the chlorophyll in suspended cells, in contrast to PilB::Tn5, which remained planktonic at this time point (Fig. 2C, day 2). By day 3, SigF::Mu showed robust biofilms, with less than 5% of the chlorophyll in suspended cells, whereas PilB::Tn5 initiated biofilm formation at this time point (Fig. 2C, day 3). Both mutants exhibited robust biofilms on day 6; however, significant differences were observed (percentage of chlorophyll in planktonic cells was 3.8%±1.1 for PilB::Tn5 mutant and 1.1%±0.8 for SigF1::Mu mutant (p<5E^-5^), Fig. 2C, day 6).

To validate the requirement of SigF1 for biofilm suppression and exclude potential polar effects on neighboring genes, the *sigF1* gene was introduced into a neutral site in the chromosome of SigF1::Mu (Table S1). The resulting strain, SigF1::Mu/F1, grew planktonically, confirming the role of SigF1 in suppressing biofilm formation.

*S. elongatus* possesses an additional *sigF* paralog (Fig. 2A), denoted *sigF2* [32], which is homologous to SigJ of *Anabaena* sp. PCC 7120 [9, 33] . This SigF variant shares the R2 and R4 domains with SigF1 but lacks the R3 domain (Fig. 2A). Deletion of *sigF2* (ΔSigF2), however, did not affect planktonic growth (Fig. 2C, day 6). This outcome indicates that SigF2 is not implicated in biofilm inhibition.

Examining the total chlorophyll accumulation up to day 6, it was found that the SigF1::Mu mutant had an approximate reduction of 25% compared to the wild-type, while no significant distinctions were observed between the WT, PilB::Tn5, and ΔSigF2 mutants (see Fig. S1). Inactivation of the single *sigF* gene of *Synechocystis* PCC 6803 also led to diminished chlorophyll accumulation [20]. Hence, SigF in this cyanobacterium as well as SigF1 in *S. elongatus* control genes related to photosynthesis.

In summary, the divergent behaviors of SigF1::Mu biofilm formation in contrast to the planktonic growth of ΔSigF2, along with the presence of the R3 domain solely in SigF1, indicate distinct cellular roles for these two sigma F factors in *S. elongatus*.

### Conditioned medium from WT cultures inhibits biofilm formation by SigF1::Mu

We previously showed that conditioned medium (CM) from WT culture effectively inhibits biofilm formation by PilB::Tn5 (Fig. 3, [22, 24]); thus, this mutant can recognize and react to a biofilm inhibitor produced and secreted by the WT strain. To investigate whether SigF1::Mu responds to or remains unaffected by externally supplied biofilm inhibitor, we introduced this mutant into WT-CM. Similar to PilB::Tn5, biofilm formation by SigF1::Mu was entirely inhibited by WT-CM (Fig. 3). In cultures of both mutants, all chlorophyll remained in the planktonic fraction, in stark contrast to the robust biofilms formed in fresh medium (Fig. 3). We conclude that SigF1::Mu, like PilB::Tn5, is impaired in synthesis or secretion of the biofilm inhibitor but is capable of sensing and responding to inhibitor that is present in WT-CM. Thus, SigF1 is not necessary for the mechanism responsible for detecting or transmitting signals from the extracellular inhibitor.

**Figure 3:**
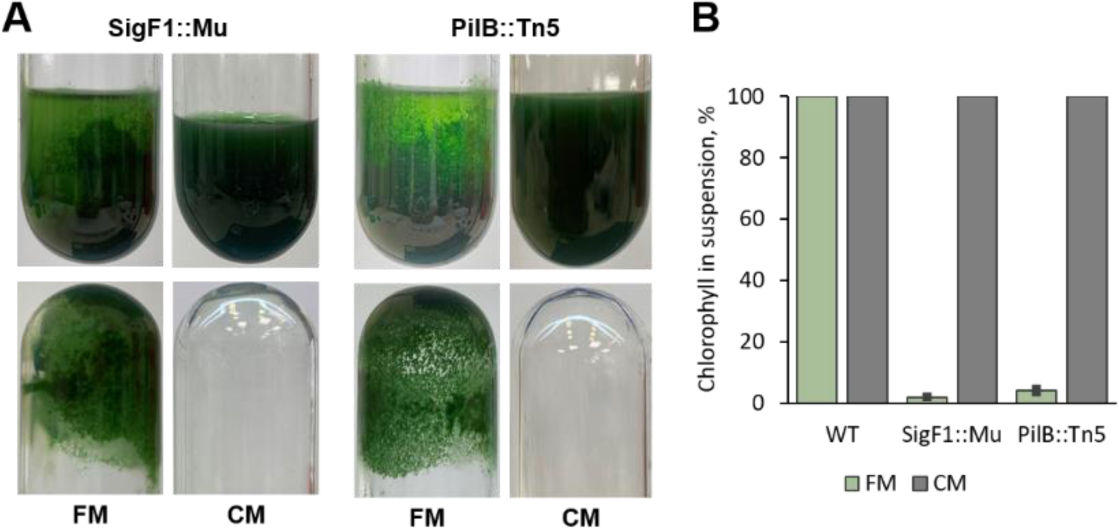
Conditioned medium from WT culture inhibits biofilm formation by SigF1::Mu. **A.** Cultures of SigF1::Mu and PilB::Tn5 in fresh medium (FM) and in WT conditioned medium (CM). Growth tubes were photographed before (upper panels) and after (lower panels) planktonic cells were decanted. In FM cells adhere to the growth vessel. **B.** Percentage of chlorophyll in suspended cells served to quantify biofilm formation.

### sigF1 shares high cofitness scores with genes encoding T4P components

To gain insight into the cellular pathways that the SigF proteins of *S. elongatus* regulate, we examined co-fitness values, which associate the loci of an RB-TnSeq library of mutants that respond similarly under a standardized set of nutritional and stress conditions [34, 35]. Genes are defined to have strong co-fitness, and potentially act in similar pathways, if they have co-fitness scores >0.75. *sigF2* mutants do not show strong co-fitness with other genes (Table S2). Inactivation of *sigF1*, however, affected fitness similarly to inactivation of genes encoding components of T4P and the DNA competence machinery, all of which have very high co-fitness values (0.89–0.99, Table 1). Additionally, mutants in genes *ebsA* and *hfq*, which encode components of the tripartite complex with PilB and are known to be essential for biofilm suppression (Fig. 1, [28]), exhibit high co-fitness with *sigF1* mutants (Table 1). These findings strongly suggest involvement of SigF1 in cellular processes associated with the T4P complex.

**Table 1:**
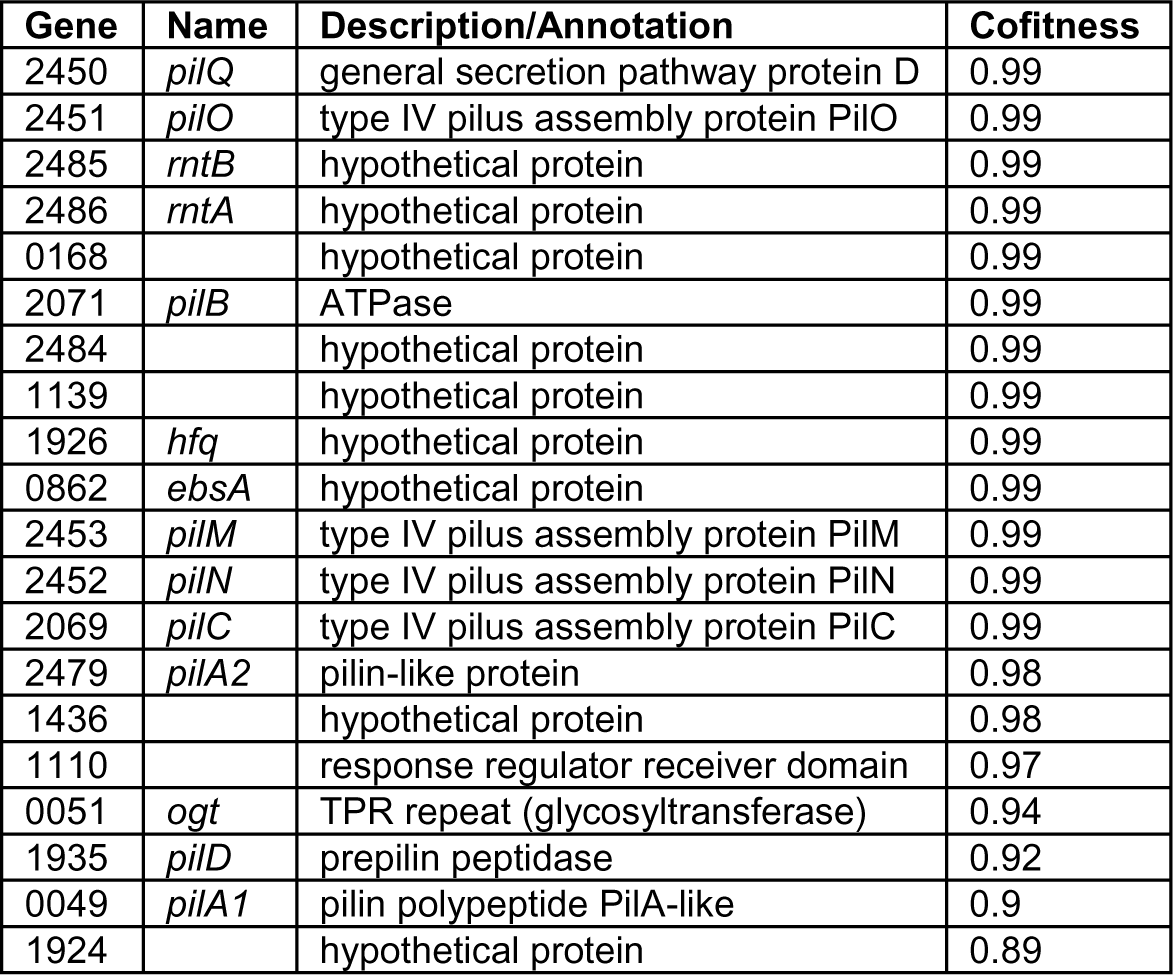
Cofitness data for *sigF1* mutants. Column ‘Gene’ describes the last four digits of full gene name (synpcc7942_XXXX). Genes Synpcc7942_2485 and Synpcc7942_2486 are required for natural transformation (*rntA* and *rntB*, [32]).

| Gene | Name | Description/Annotation | Cofitness |
| --- | --- | --- | --- |
| 2450 | *pilQ* | general secretion pathway protein D | 0.99 |
| 2451 | *pilO* | type IV pilus assembly protein PilO | 0.99 |
| 2485 | *rntB* | hypothetical protein | 0.99 |
| 2486 | *rntA* | hypothetical protein | 0.99 |
| 0168 |  | hypothetical protein | 0.99 |
| 2071 | *pilB* | ATPase | 0.99 |
| 2484 |  | hypothetical protein | 0.99 |
| 1139 |  | hypothetical protein | 0.99 |
| 1926 | *hfq* | hypothetical protein | 0.99 |
| 0862 | *ebsA* | hypothetical protein | 0.99 |
| 2453 | *pilM* | type IV pilus assembly protein PilM | 0.99 |
| 2452 | *pilN* | type IV pilus assembly protein PilN | 0.99 |
| 2069 | *pilC* | type IV pilus assembly protein PilC | 0.99 |
| 2479 | *pilA2* | pilin-like protein | 0.98 |
| 1436 |  | hypothetical protein | 0.98 |
| 1110 |  | response regulator receiver domain | 0.97 |
| 0051 | *ogt* | TPR repeat (glycosyltransferase) | 0.94 |
| 1935 | *pilD* | prepilin peptidase | 0.92 |
| 0049 | *pilA1* | pilin polypeptide PilA-like | 0.9 |
| 1924 |  | hypothetical protein | 0.89 |

### Transcriptome analyses of SigF1::Mu, ΔSigF2 and PilB::Tn5

To study the role of SigF1 in transcriptional regulation, we conducted comparative analyses of the global transcriptome between the WT strain and SigF1::Mu mutant. PilB::Tn5 was also included in these analyses to help identify biofilm-related transcriptional changes. Additionally, we examined ΔSigF2 to understand how the transcriptional space is divided between the two sigma F factors of *S. elongatus*. RNA extraction was performed at two time points: day 1, when all strains were in planktonic growth, and day 4, when both PilB::Tn5 and SigF1::Mu had developed biofilms. Venn diagrams summarize differentially expressed genes (DEGs) in the mutants relative to WT (Fold > 2, adjusted p value <0.05, Fig. 4 and Supplemental file 1).

**Figure 4:**
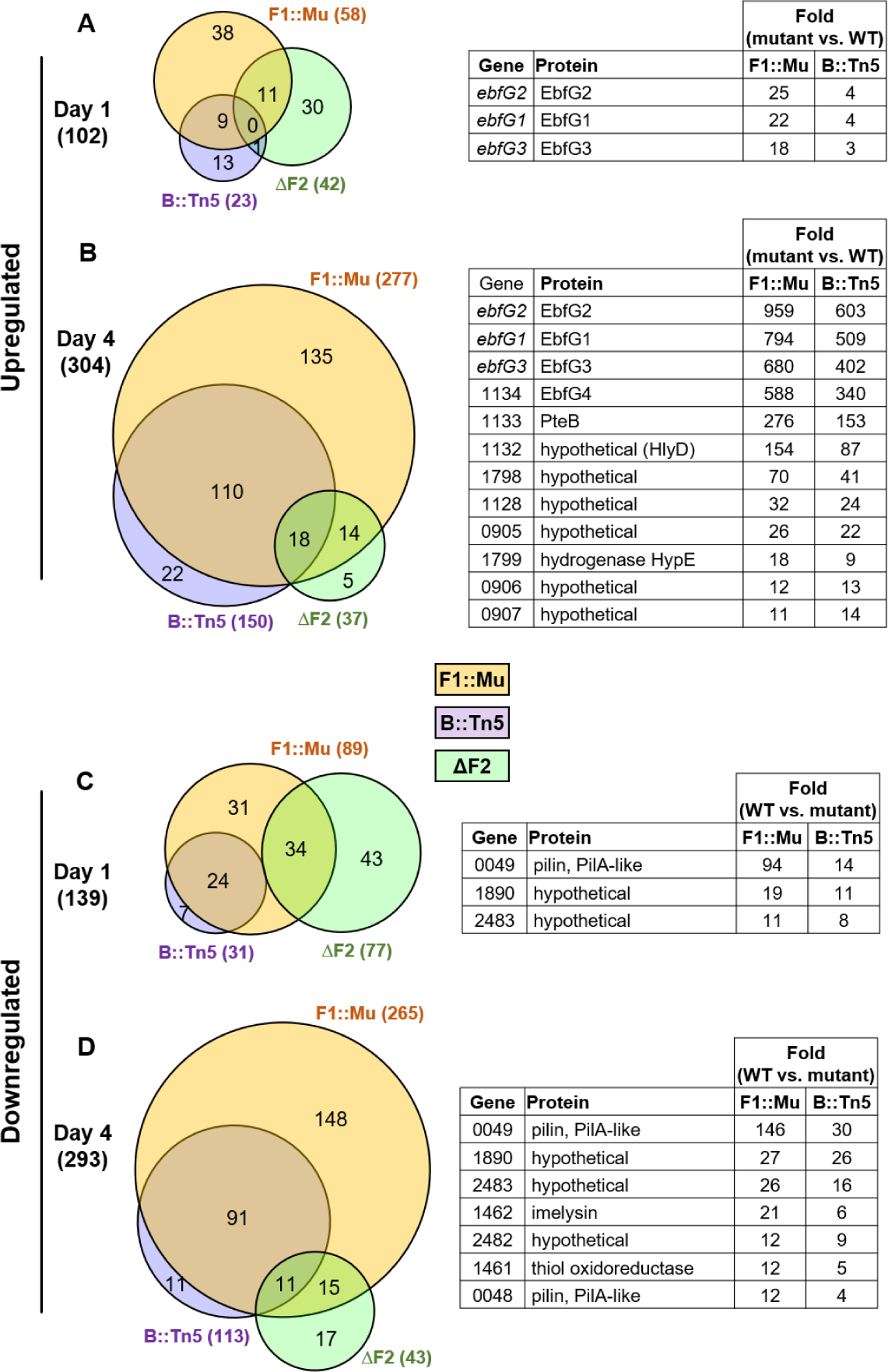
Summary of transcriptome analyses of WT, PilB::Tn5 (B::Tn5,) SigF1::Mu (F1::Mu) and ΔSigF2 (ΔF2). Venn diagrams summarize differentially expressed genes (DEGs) that were up- or downregulated in the mutants relative to WT (fold change ≥ 2; adjusted p < 0.05). Tables indicate protein encoding genes exhibiting the highest fold change for SigF1::Mu compared to WT (fold change > 10; adjusted p < 0.05), which are also differentially expressed in PilB::Tn5 but not in ΔSigF2. For complete data set see Supplementary File S1. **A** and **B**. Upregulated genes in mutants compared to WT in day 1 or day 4, respectively. **C** and **D**. Downregulated genes in mutants compared to WT in day 1 or day 4, respectively.

#### Distinct regulons of SigF1 and SigF2

Transcriptome analyses of 4-day old cultures revealed substantially different regulons of SigF1 and SigF2. Only approximately 10% of the genes up- or downregulated in SigF1::Mu relative to WT were shared with ΔSigF2 (32 of 277, and 26 of 265, respectively). Notably, at this time point, SigF1 governs a larger regulon compared with SigF2 (Fig. 4 B&D; 277 vs. 37 upregulated genes and 265 vs. 43 downregulated genes). The observed differences in regulons between SigF1 and SigF2 can likely be attributed to the distinct domain compositions of these sigma factors, which may affect their promoter specificity and consequently dictate the regulation of different sets of genes. Moreover, the presence of both up- and downregulated genes in each mutant suggests that both SigF factors are involved in transcriptional repression of specific gene sets and activation of other gene sets. Upregulation of genes in SigF1::Mu implies that, in the absence of SigF1, another sigma factor or response regulator activates transcription of this set of genes. Possibly, the putative positive transcriptional activator drives transcription in WT cells when SigF1-modulation, possibly by posttranslational modification or anti-sigma factors, alleviates its repressive effect. Cyanobacterial anti-sigma factor candidates have been predicted based on sequence similarity to known factors from heterotrophic bacteria; however, most have not been verified experimentally [1]. A study of *Synechocystis* PCC 6803 demonstrated that the H subunit of Mg-chelatase serves as a SigE-anti-sigma factor, thereby broadening the scope of proteins that may serve for sigma factors regulation [36]. An additional study implicated SapG, a SigG anti-sigma factor, in regulation of envelope stress in *Nostoc punctiforme* [37]. In addition, one may envision a regulatory scenario in which SigF1 represses transcription of a positive transcription activator.

#### High transcript levels of biofilm-promoting genes in SigF1::Mu

Transcriptome analyses revealed substantial upregulation of the *ebfG*-operon in SigF1::Mu. The transcript abundances of *ebfG* genes were 18-25 fold higher on day 1 and ∼600-1000 fold higher on day 4 in SigF1::Mu compared to WT (Fig. 4 A&B and Supplemental file 1). Additionally, *pteB*, which encodes a cysteine peptidase that takes part in secretion and maturation of EbfG proteins [24], is upregulated approximately 270-fold in SigF1::Mu compared to WT (Fig. 4B).

Venn diagrams in Fig. 4B-D show that most DEGs of PilB::Tn5 are encompassed within the DEGs of SigF1::Mu. Possibly, these shared transcriptional changes between the two biofilm-forming strains represent processes related to biofilm formation or cellular functions associated with proliferation or survival within the biofilm environment. One of these genes, synpcc7942_1132, which encodes a homolog of HlyD, a component of type I secretion systems, is upregulated in the biofilm-forming mutants (Fig. 4B). Given involvement of a type I system in secretion of the EbfG proteins [24, 26] we investigated the role of *hlyD* in biofilm formation by constructing the double mutant PilB::Tn5/HylD::Mu (Table S1). This strain grew planktonically (Fig. S2); therefore, we concluded that HlyD is required for biofilm formation, most likely due to its involvement in secretion of the EbfG proteins.

Notably, on day 1, higher levels of transcripts of *ebfG*1-3 were observed in SigF1::Mu compared to PilB::Tn5 (Fig. 4A), consistent with the earlier biofilm formation manifested by SigF1::Mu (Fig. 2C). Additionally, a larger increase in transcript levels of the *ebfG*-operon and the biofilm-related genes *pteB* and *hlyD* was observed in SigF1::Mu compared to PilB::Tn5 (Fig. 4 A&B).

#### Low transcript levels of *pilA1* genes in SigF1::Mu

The major pilus subunit in *S. elongatus* is encoded by two adjacent genes, synpcc7942_0049 and synpcc7942_0048 [27, 30, 32], both referred to as PilA1 due to their 98% amino acid identity. The majority of *pilA1* transcripts in WT cells originate from gene synpcc7942_0049 (Fig. S3). Transcript levels of this gene are 94-fold higher on day 1 and 146-fold higher on day 4 in WT compared to SigF1::Mu (Fig. 4C&D; see also Supplemental file 1). Similar, although less pronounced, differences are observed for synpcc7942_0048 (Fig. 4D). These findings suggest that SigF1 plays a role in activating the transcription of the *pilA1* genes, which, in turn, affects the secretion of the biofilm inhibitor. Of note, SigF of *Synechocystis* has been implicated in transcription activation of *pilA1* [17, 19].

Gene Synpcc7942_2482 that encodes a PilA candidate [38] is downregulated in SigF1::Mu and PilB::Tn5. Inactivation of this gene, however, resulted in planktonic growth [27] and thus, this putative pilin subunit is not required for secretion of the biofilm inhibitor. Functional redundancy may also explain the observed phenotype.

#### Impact of sigF1 inactivation on pili formation and DNA competence

Given that *pilA1* transcript is substantially less abundant in SigF1::Mu compared to WT, we examined piliation of this mutant by transmission electron microscopy (TEM). Pili were not apparent in SigF1::Mu (Fig. 5C) in contrast to piliated WT cells (Fig. 5A). Moreover, detached pili observed in WT cultures (Fig. 5B), were not detected in SigF1::Mu cultures. Together, low abundance of *pilA1* transcripts in SigF1::Mu compared to WT (Fig. 4 C&D) and absence of pili in this mutant (Fig. 5) are in-line with the high co-fitness of *sigF1* mutants with mutants in components of the T4P complex (Table 1). In contrast to SigF1::Mu, the ΔSigF2 mutant is piliated (Fig. 5D), in agreement with normal transcription of the *pilA1* genes in this strain (Supplemental file 1).

**Figure 5:**
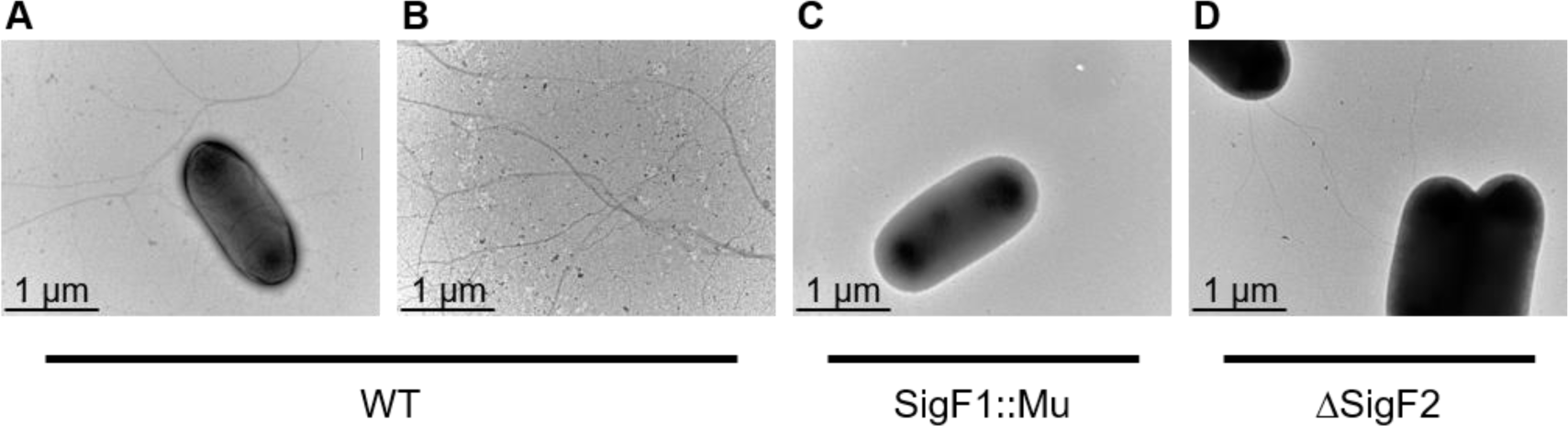
Transmission electron microscopy (TEM) analyses of WT, SigF1::Mu and ΔSigF2. **A, C** and **D.** Cell images. **B.** Detached cell pili observed in WT culture.

SigF involvement in transcription regulation of *pilA* and pilus formation was demonstrated in *Synechocystis* PCC 6803 and *Nostoc punctiforme* [13, 19, 20, 39]. Together with current observations of impaired *pilA1* transcription and absence of pili in SigF1::Mu of *S. elongatus*, data suggest that SigF involvement in pilus formation is a general cyanobacterial mechanism.

The T4P complex is typically associated with uptake of external DNA [32, 40–44]. Furthermore, inactivation of *sigF* in *Synechocystis* PCC 6803 abolishes DNA competence [17, 18]. Comparison of DNA competence in SigF1::Mu and ΔSigF2 revealed that SigF1::Mu is characterized by DNA competence similar to WT whereas ΔSigF2 is non-transformable (Fig. S4), in accordance with a genetic screen for genes required for DNA competence [32]. These data, together with the TEM (Fig. 5) and the transcript analyses (Fig. 4C&D), suggest that pili comprising PilA1 are those observed in TEM images of WT and ΔSigF2. These pili, which are absent from SigF1::Mu are dispensable for DNA competence. Probably, the SigF1::Mu strain assembles few, thin, or short pili that are not observed by the negative staining and TEM analysis and that are functional in DNA uptake. Additionally, a glycosyltransferase mutant of S. elongatus, which does not glycosylate PilA1 and the majority of its cells are non-piliated, is nevertheless transformable similar to WT [30]. *S. elongatus* possesses several alternative pilin candidate genes [27, 32, 38]; however, the identity of the competence pili is, as yet, unknown.

### EbfG expression examined by flow cytometry

Following up on the transcriptome results, which indicate that SigF1 is required for *ebfG*-operon transcription upregulation, we examined expression of this operon in individual cells using reporter strains and flow cytometry. The reporter construct comprises the putative promoter region of the *ebfG*-operon along with the 5’ untranslated region attached to a yellow fluorescence protein (*yfp*) gene, yielding the construct P-*ebfG*::YFP. This reporter gene was inserted in a neutral site in the chromosome in WT, PilB::Tn5, SigF1::Mu and ΔSigF2 cells, yielding their cognate reporter strains (Table S1 and [25]).

WT-reporter 2-day-old cultures served for defining YFP-positive cells for all reporter strains (dashed line in Fig. 6A, also see Fig. S5B). Analysis of 2-day-old cultures, prior to biofilm development by strains capable of forming biofilms, did not exhibit significant changes between WT-, PilB::Tn5- and SigF1::Mu-reporter strains (Fig. 6C). Following biofilm development by PilB::Tn5- and SigF1::Mu-reporter cultures (day 4 and day 6), biofilm and planktonic fractions were separated and measured individually (see Methods). Initial analysis, however, did not indicate significant changes between percentage of YFP-positive cells in the biofilm or the planktonic fractions of a particular mutant strain, in agreement with previous analyses of a PilB::Tn5-reporter [25]; therefore, data from the two fractions were analyzed collectively. Mutant reporters were distinct from the WT-reporter strain in both 4-day and 6-day old cultures (Fig. 6C).

**Figure 6:**
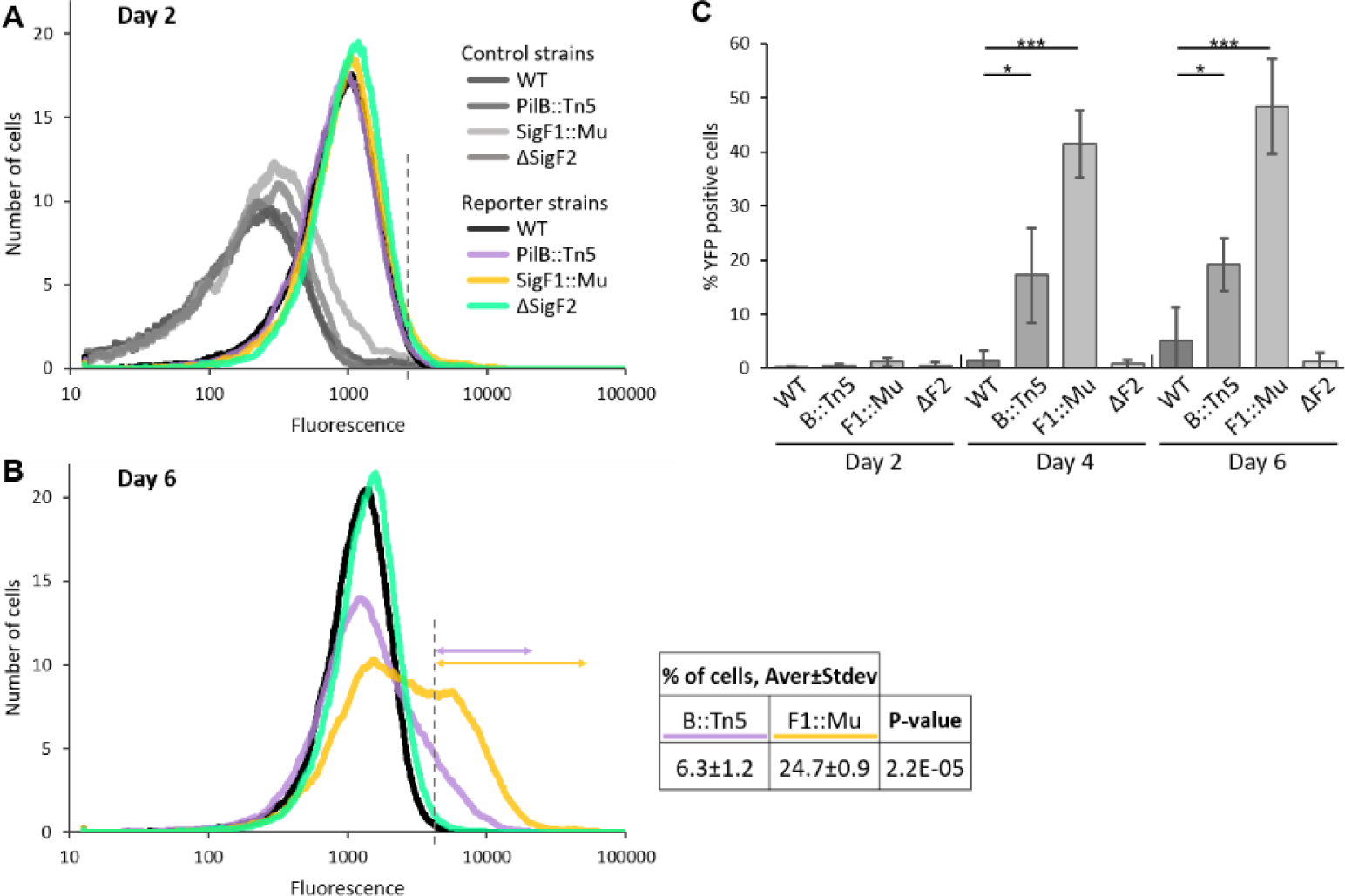
Examination of expression of the *ebfG*-operon using reporter strains and flow cytometry. **A** and **B.** Number of cells as a function of fluorescence in WT, PilB::Tn5, SigF1::Mu and ΔSigF2 strains bearing a reporter construct (P-ebfG::YFP). Analyses were performed on 2-day **(A)** and 6-day **(B)** old cultures. In **A** the cognate negative control strains are also included. Histograms depicting data from a single representative experiment out of the three biological replicates are presented. Dashed line in **A** indicates cutoff for calculating YFP positive cells (data summary in **C**). Additional cutoff (dashed line in **B**) served for calculating YFP positive mutant-reporter cells relative to 6-day old WT-reporter (see Table for averages and standard deviations from three biological replicates). **C.** Percentage of YFP-positive cells in PilB::Tn5 (B::Tn5), SigF1::Mu (F1::Mu) and ΔSigF2 (ΔF2) reporter cultures relative to 2-day old WT-reporter cells. Averages and standard deviations from three biological replicates are presented. Asterisks denote significant changes between WT- and mutant-reporter strains of same culture age (one-way ANOVA with Dunnett’s post-hoc test. * p<0.05; *** p<0.001).

In addition to using WT-reporter 2-day-old cultures for defining YFP-positive cells (data shown in Fig. 6C), we used 6-day-old WT-reporter cultures to define YFP-positive cells in PilB::Tn5 and SigF1::Mu reporter strains at this culture stage (Fig. 6B, dashed line). This analysis revealed approximately 25% YFP-positive cells in the SigF1::Mu-reporter compared to around 6% in the PilB::Tn5-reporter (Fig. 6B).

Notably, only a subfraction of cells in cultures of the biofilm-forming strains exhibits high expression of YFP (Fig. 6B&C), indicating cell specialization in expression of the *ebfG*-operon, in agreement with previous analyses of PilB::Tn5 [25]. EbfG4 is a surface and matrix-localized protein and EbfG1-3, which are prone to form amyloids, are likely to be matrix components [25]. The SigF1::Mu data support the premise that, through cell specialization in expression of the *ebfG*-operon, production of “public goods” by part of the population is sufficient to support biofilm development by the majority of the cells. Moreover, the higher fraction of YFP-positive cells in SigF1::Mu compared to PilB::Tn5 (Fig. 6B) is in-line with the more vigorous biofilm produced by SigF1::Mu (Fig. 2C, day 6).

Expression of the *ebfG*-operon in ΔSigF2 did not significantly differ from WT throughout the experiment (Fig. 6), in agreement with the transcriptome analyses (Supplemental file 1) and the planktonic nature of this mutant (Fig. 2).

### Exoproteome analyses reveal substantially higher level of EbfG proteins in SigF1::Mu than in PilB::Tn5

Our previous exoproteome analyses indicated higher amounts of EbfG proteins in CM of PilB::Tn5 compared to WT [27]. Elevated expression of the *ebfG*-operon and components of the type-I secretion system related to secretion of the EbfG proteins in SigF1::Mu compared to PilB::Tn5 encouraged us to compare the exoproteomes of these two mutants. Data revealed 50-3400-fold higher extracellular level of EbfG proteins in SigF1::Mu compared to PilB::Tn5 (Figure 7A). This finding is in-line with observed earlier onset of biofilm formation SigF1::Mu compared to PilB::Tn5 (Fig. 2C).

**Figure 7:**
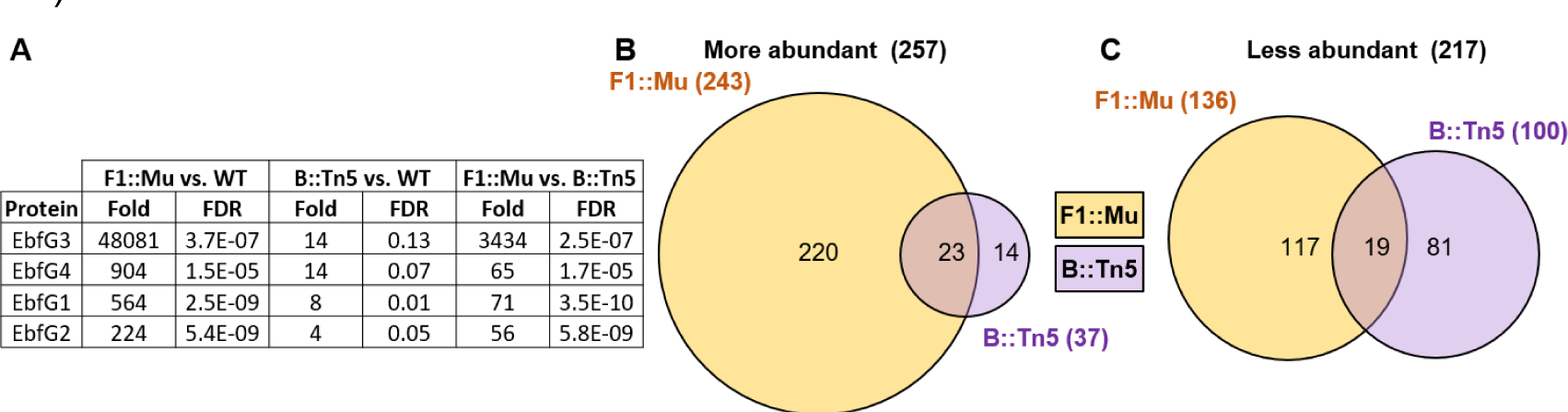
Exoproteome analyses reveal substantially higher level of EbfG proteins in SigF1::Mu than in PilB::Tn5. **A.** Table summarizing enrichment of EbfG proteins in mutant exoproteomes. Venn diagrams summarizing more (**A**) and less (**B**) abundant proteins in the exoproteomes of SigF1::Mu and PilB::Tn5 compared to WT (fold change > 2; FDR <0.1).

Additionally, the exoproteome of SigF1::Mu is characterized by numerous proteins that are more or less abundant compared to WT (Fig. 7 B&C). These changes likely represent SigF1 impact on different secretion processes. Inactivation of *sigF* in *Synechocystis* PCC 6803 also substantially affected the exoproteome [20]. Together, these observations imply that regulation of secretion processes by SigF is a common cyanobacterial mechanism.

Genes synpcc7942_0905 and synpcc7942_0906 whose transcripts are highly elevated in SigF1::Mu and PilB::Tn5 compared to WT (Fig. 4B) encode proteins of unknown function or annotation that are more abundant in the exoproteome of these mutants compared to WT (Supplemental file 2; independent exoproteome analysis of PilB::Tn5 also indicated enrichment of these proteins compared to WT [27]). Given elevated expression of these genes, and their encoded proteins in the exoproteomes of the biofilm-forming mutants, we proposed that, similarly to products of the *ebfG*-operon, these proteins may contribute to matrix formation. To test this hypothesis, a double mutant in which *pilB* was inactivated along with deletion of synpcc7942_0905 and synpcc7942_0906 was constructed. This strain, however, demonstrated vigorous biofilm formation, comparable to PilB::Tn5 (Fig. S6), negating involvement of these gene products in biofilm formation.

### Summary model: SigF1 regulates biofilm formation via intra- and intercellular pathways

The data strongly support the involvement of SigF1 in the intricate regulation of genes that promote biofilm formation, acting through two distinct pathways as illustrated in Figure 8. This alternative sigma factor plays a fundamental role in activation of *pilA1* transcription as evident by the substantially low transcript levels of the major pilin subunit (Fig. 4 C&D) in SigF1::Mu and consequently, the major pili are absent from this mutant (Fig. 5). The T4P complex has been assigned a role in deposition of biofilm inhibitor(s) to the extracellular milieu. This action subsequently leads to the repression of genes that promote biofilm formation, which in turn represses biofilm-promoting genes (Fig. 8, top panel, thick T-bar; [21–25]). Therefore, *sigF1*-inactivation, which abrogates pili formation, alleviates a major repression pathway, most likely due to disruption of the inhibitor(s) secretion process. This conclusion is supported by planktonic growth of SigF1::Mu when inoculated into WT-CM (Fig. 2). Additionally, the SigF1::Mu response to inhibitor(s) within WT-CM underscores that SigF1 involvement is not essential for the perception or transduction of the inhibitory signal.

**Figure 8:**
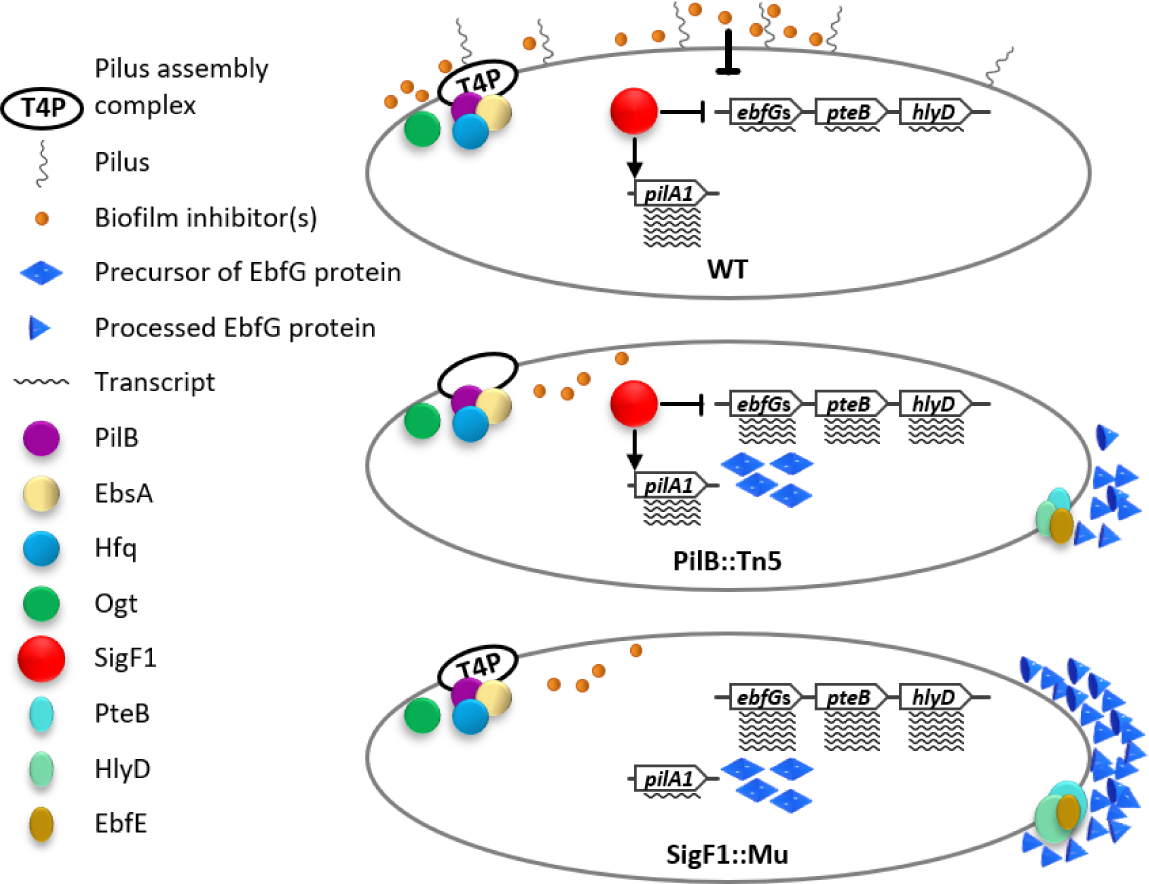
Dual repression pathways by SigF1 in biofilm regulation. SigF1 regulates transcription of biofilm-promoting genes via an intracellular mechanism (thin T-bar) as well as intercellular pathway (thick T-bar). See text for further details.

Furthermore, we propose that SigF1 is involved in an additional inhibitory pathway, based on the higher expression of biofilm-promoting genes in SigF::Mu compared to PilB::Tn5 as supported by three lines of experiments: Higher levels of transcripts from the *ebfG*-operon, *pteB* and *hlyD* (Fig. 4) and elevated EbfG expression that is manifested by flow cytometry data (Fig. 6) and exoproteome analyses (Fig. 7). In PilB::Tn5, where the assembly ATPase of the T4P complex is inactivated, the repression exerted by the extracellular inhibitor is nullified; however, SigF1 contributes to repression of EbfGs via an intracellular pathway (Fig. 8, middle panel, thin T-bar). In SigF1::Mu this pathway is abolished along with the intercellular pathway mediated by secreted inhibitor(s); consequently, transcription repression in SigF1::Mu is completely alleviated (Fig. 8). The ubiquitous presence of EbfG proteins in the SigF1::Mu exoproteome results from the remarkably high expression of the *ebfG*-operon and the related secretion machinery (Figs. 7&8).

EbfG4 functions as both a cell-surface and matrix protein, and EbfG1-3 exhibit a propensity to form amyloids [25]. These observations, coupled with findings from studies in heterotrophic bacteria that attribute a role to amyloids as fundamental building blocks of the matrix [45–48], collectively support the robust matrix formation potential of SigF1::Mu.

Accumulation of biofilm inhibitor with culture growth and increase in cell density support a quorum-like mechanism in regulation of *S. elongatus* biofilm formation [25]. Of note, the field of cyanobacterial quorum sensing and regulation of intercellular signaling is in its infancy. Thus, this study, which assigns a role to SigF1 in intercellular communication, provides a step forward in elucidation of these cyanobacterial mechanisms.

The *Synechocystis sigF* mutant exhibited clumping and increased polysaccharide secretion: phenotypes that are often associated with biofilm formation. However, biofilms were not reported for this mutant [20]. It is conceivable that under different growth conditions, this mutant could develop biofilms and, therefore, we propose that SigF involvement in biofilm regulation is likely a more widespread characteristic shared among cyanobacteria.

## Materials and Methods

### Strains, culture conditions, biofilm assay and competence analysis

*S. elongatus* PCC 7942, an obligatory photoautotroph, and all derived strains were grown in mineral medium BG11 as described [31]. Cultures were grown at 30°C in Pyrex tubes under bubbling with air enriched with CO_2_. Incandescent light was provided at flux of ∼30 μmol photons m^−2^ sec^−1^. Construction of mutants and details of molecular manipulations are provided in Supplementary Table 1.

Biofilm quantification in bubbled cultures is based on chlorophyll measurement as a proxy for biomass accumulation in sessile as well as in planktonic cells and representation of the relative fraction of chlorophyll in planktonic cells. Chlorophyll was extracted in 80% acetone quantified based on absorbance at optical density at 663 nm [31]. Biofilms formed under static conditions in 24-well plates at 28°C with incandescent light illumination (30 μmol photons m^-2^ s^-1^) were quantified by crystal violet stanning following 9 days essentially as described [49] except for crystal violet extraction that was performed in 95% ethanol and not in 30% acetic acid.

Assessment of DNA competence was performed essentially as described previously[50]. Exponentially growing cells were centrifuged (5000g for 8 min at room temperature), washed once with 10 mM NaCl, and re-suspended to an OD_750_ of 4.0. A shuttle vector (1000 ng) was added to 600 μl of cells, which were gently agitated overnight at 28°C in the dark. Transformants were selected by plating on selective solid growth medium (50 μg ml^-1^ spectinomycin) supplemented with NaHCO_3_ (5 mM) and sodium thiosulfate (0.3%, wt./vol.). The shuttle vector replicates autonomously, thus allowing the assessment of DNA uptake without a possible impact on the efficiency of DNA integration into the chromosome.

### Microscopy

To observe biofilms by fluorescence microscopy, a sterile microscope slide was inserted into a growth tube upon culture inoculation, and biofilms formed on the glass tube wall as well as on the microscope slide. Following 7 days of growth, the microscope slide was removed with forceps and washed once by dipping into double-distilled water. Autofluorescence-based images were collected using a Leica SP8 confocal microscope (excitation at 630nm and emission at 641 to 657 nm)[28].

For transmission electron microscopy, one day old cultures that had not yet initiated biofilm formation were sampled (10 μl) and applied onto ultra-thin carbon-coated grids. Following 5 min liquid was removed by blotting and cells were negatively stained twice with 2% fresh aquatic uranyl acetate (10 μl) for 1 min each step. Stain was removed by blotting, grids were briefly washed with double distilled water (10 μl) and left to dry overnight at room temperature. Images were acquired with a Tecnai G2 Fei transmission electron microscope, operating at 120 kV with a 1KX1K camera[30].

### RNA extraction and transcriptome analysis

Four biological repeats were grown under bubbling and RNA extraction was performed 1 and 4 days after inoculation. For 4-days old PilB::Tn5 and SigF1::Mu cultures, supernatant fraction was carefully removed, and the biofilm fraction was resuspended in 3 ml of remaining supernatant. Total RNA was extracted as previously described [21]. 40 µg of total RNA was treated with 4 U of TURBO™ DNase (Ambion, Catalog #: AM2238) at room temperature for 45 min followed by a boost with 4 U and an additional 45 min incubation. The DNase was removed by phenol-chloroform and chloroform extractions, RNA was precipitated and re-treated with DNAse as described above. RNA sample pellets resulting from the ethanol precipitation were washed with ice cold 70% ethanol, air-dried and resuspended in RNase-free water. 3 μg of RNA from each sample was depleted of ribosomal RNA using the RiboMinus™ Pan-Prokaryote Probe Mix, Invitrogen™ A46920. RNA quality control was performed using 1% agarose gel electrophoresis and The Agilent TapeStation system. Strand-specific library preparation and sequencing on an Illumina NextSeq system was performed on purified ribosomal RNA-depleted samples at The NGS unit, Kanbar core facility center at Bar-Ilan University.

Sequenced reads were mapped to the *S. elongatus* PCC 7942 reference genome using Rockhopper [51–53]. Normalization and differentially expressed gene test were implemented by DESeq2. An arbitrary cutoff of at least 1 log2-fold and p-value adjusted for multiple testing < 0.05 were chosen to define DEGs.

### Flow cytometry

Aliquots of 0.5 ml were taken from each culture tube following 2, 4 and 6 days of growth and then, in case of biofilm-forming strains, planktonic cells were removed. 1.5 ml BG11 were used to resuspend the biofilmed cells by rigorous pipettation and 0.13 ml were transferred to a 1.5 ml Eppendorf tube for homogenization with a pellet pestle (Sigma-Aldrich, Z359971-1EA). The homogenized samples were filtered through a mesh (pore size 52 µm), supplemented with formaldehyde to a final concentration of 1%, diluted with phosphate-buffered saline (PBS) to OD750 of ∼0.0001 and measured using BD FACSAria (excitation 488nm, emission 530 ±30nm). Gating for flow cytometry analysis was based on cyanobacterial autofluorescence (Fig. S5A).

### Exoproteome analysis

Harvesting of CM and MS analyses were performed as previously described [28]. Briefly, for collection of CM, cultures were centrifuged (5,000 x g for 10 min) at room temperature, and the supernatant was removed and passed through a 0.22 µm filter. Data were analyzed to identify proteins that are significantly more or less abundant in a particular mutant’s exoproteome than in the WT (FDR of ≤0.1), with a cutoff of at least a 2-fold change. Comparative analysis of WT- and PilB::Tn5-exoproteome was included in our previous publication [28].

## Supplementary information

**Figure S1:**
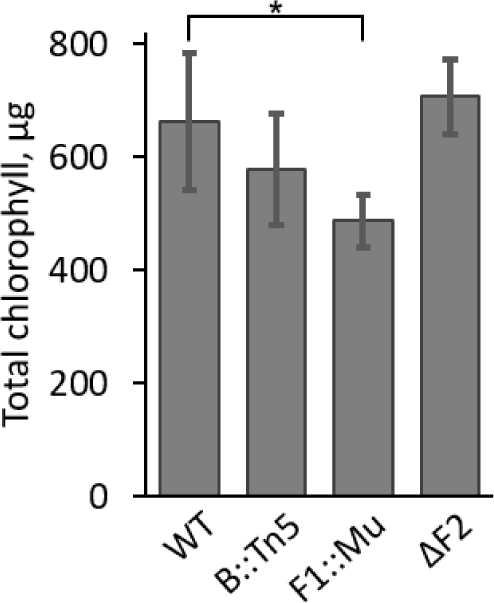
Total chlorophyll in 6-day old cultures of WT, PilB::Tn5 (B::Tn5), SigF1::Mu (F1::Mu) and ΔSigF2 (ΔF2). * F1::Mu significantly differ from WT (t-test, two tails, two-sample assuming unequal variances. p<0.00035).

**Figure S2:**
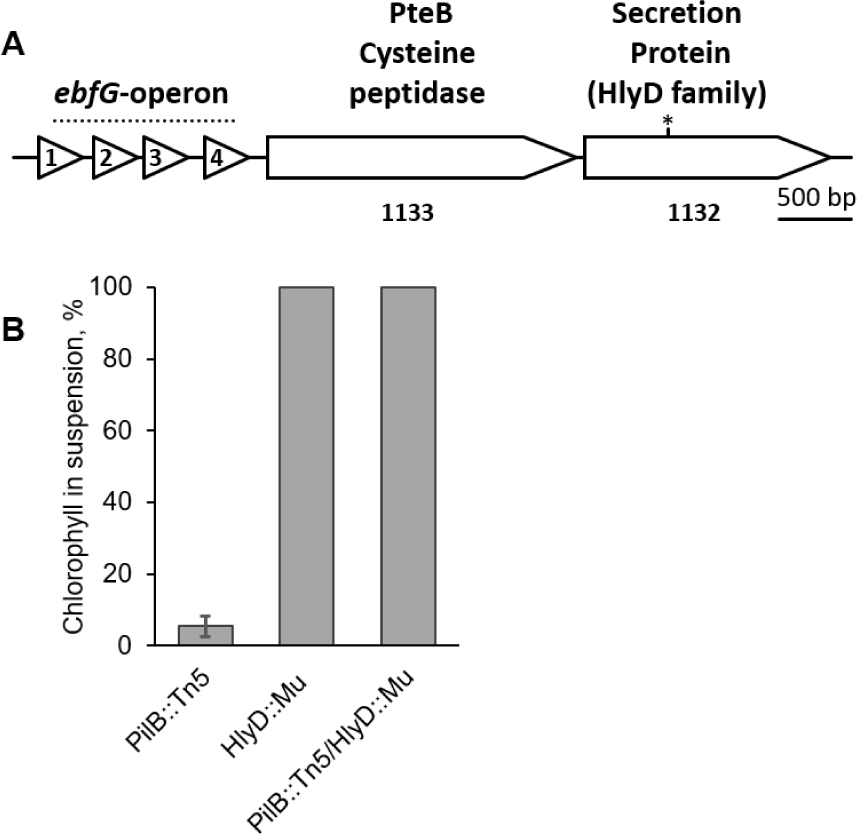
**A.** Genomic region of gene synpcc7942_1132 encoding a HlyD homolog. Asterisk indicates insertion site of Mu-transposon based cassette. **B.** Assessment of biofilm development by measurement of the percentage of chlorophyll in suspended cells. Robust biofilm development is manifested in low percentage of the chlorophyll in suspended cells. Strains analysed: PilB::Tn5; HlyD::Mu and PilB::Tn5/HlyD::Mu. Data represent averages and standard deviations from three independent experiments.

**Figure S3:**
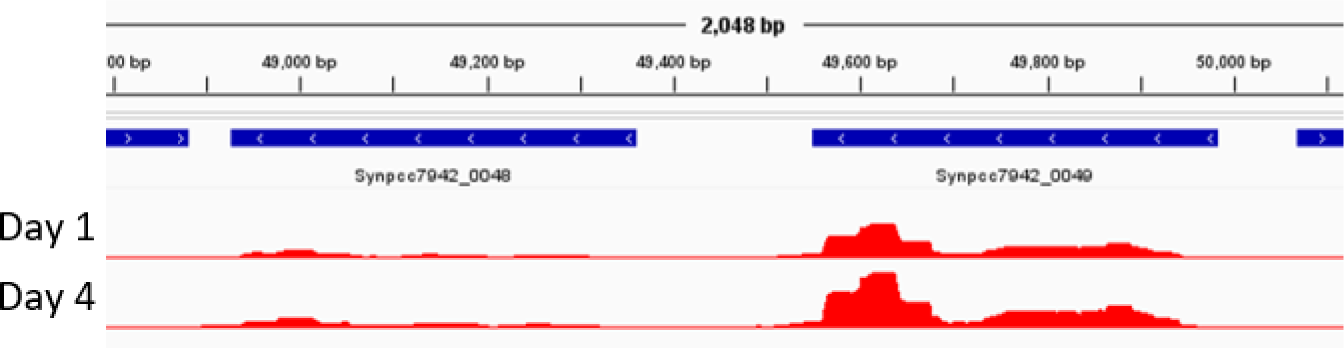
Visualization of *pilA1* transcripts in one- and four-day old cultures of WT. Transcript abundance is depicted using Rockhopper (see Methods).

**Figure S4:**
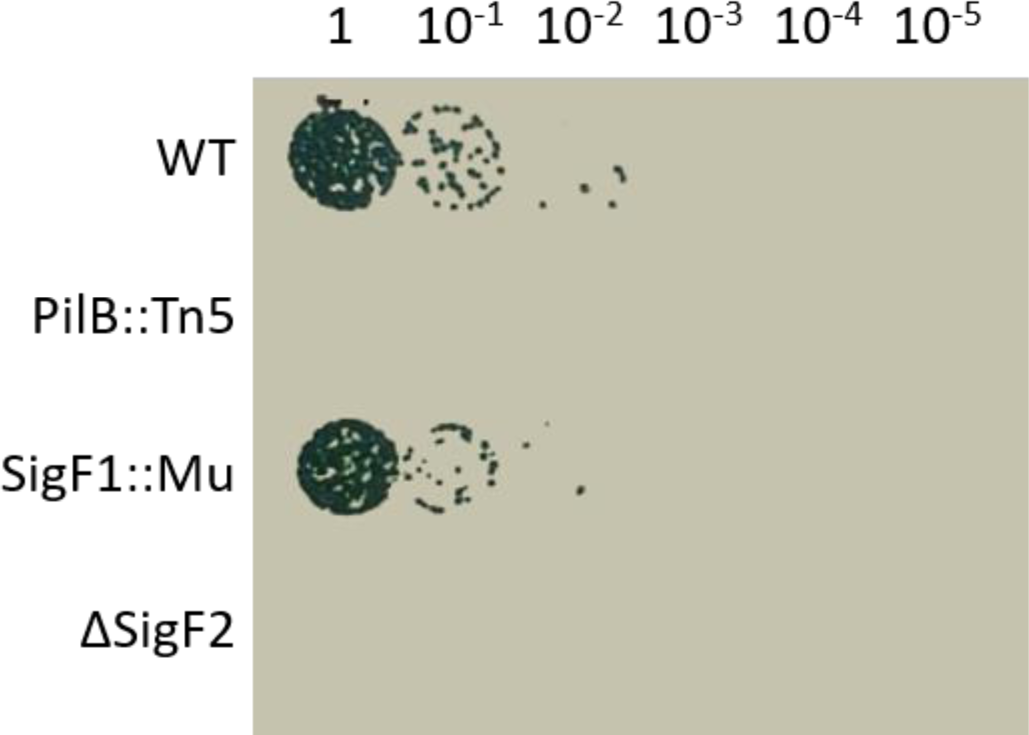
Analysis of DNA competence in WT, PilB::Tn5, SigF1:Mu and **ΔSigF2.** Shown are colonies growing on selective medium following “spotting” of 10 µl of the original transformation mixture and the indicated serial dilutions.

**Figure S5:**
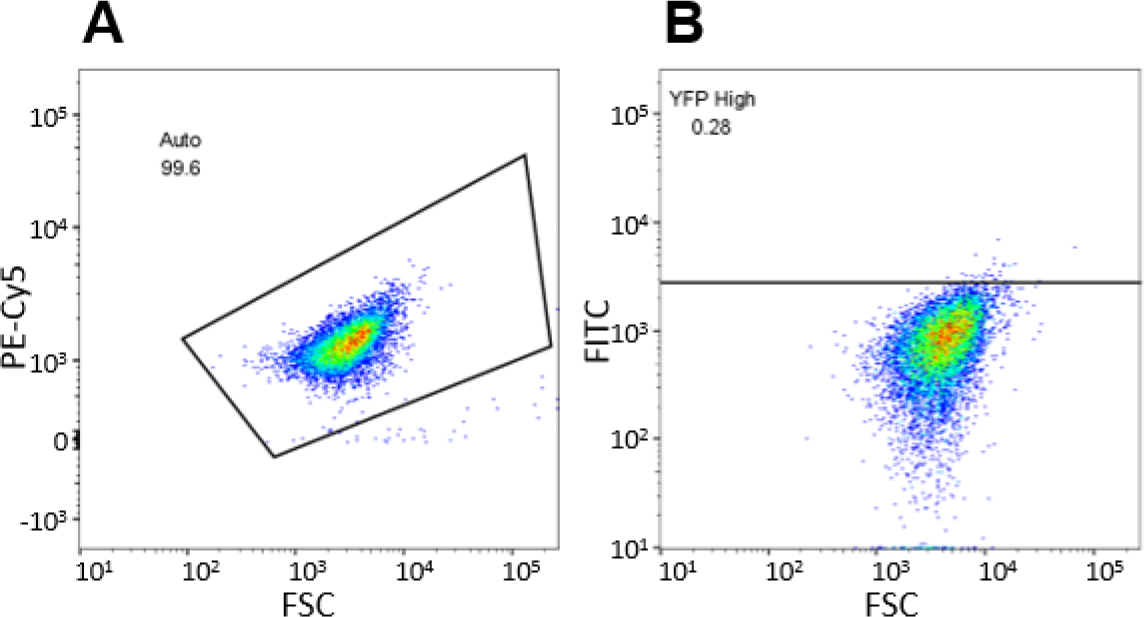
**A.** Gating for analysis of flow cytometry data is based on autofluorescence (PE-Cy5) *vs.* forward scattering (FSC). **B.** Density plots (green fluorescence (FITC *vs.* FSC) demonstrating the cutoff for calculating YFP positive cells based on WT-reporter 2-day-old cultures.

**Figure S6:**
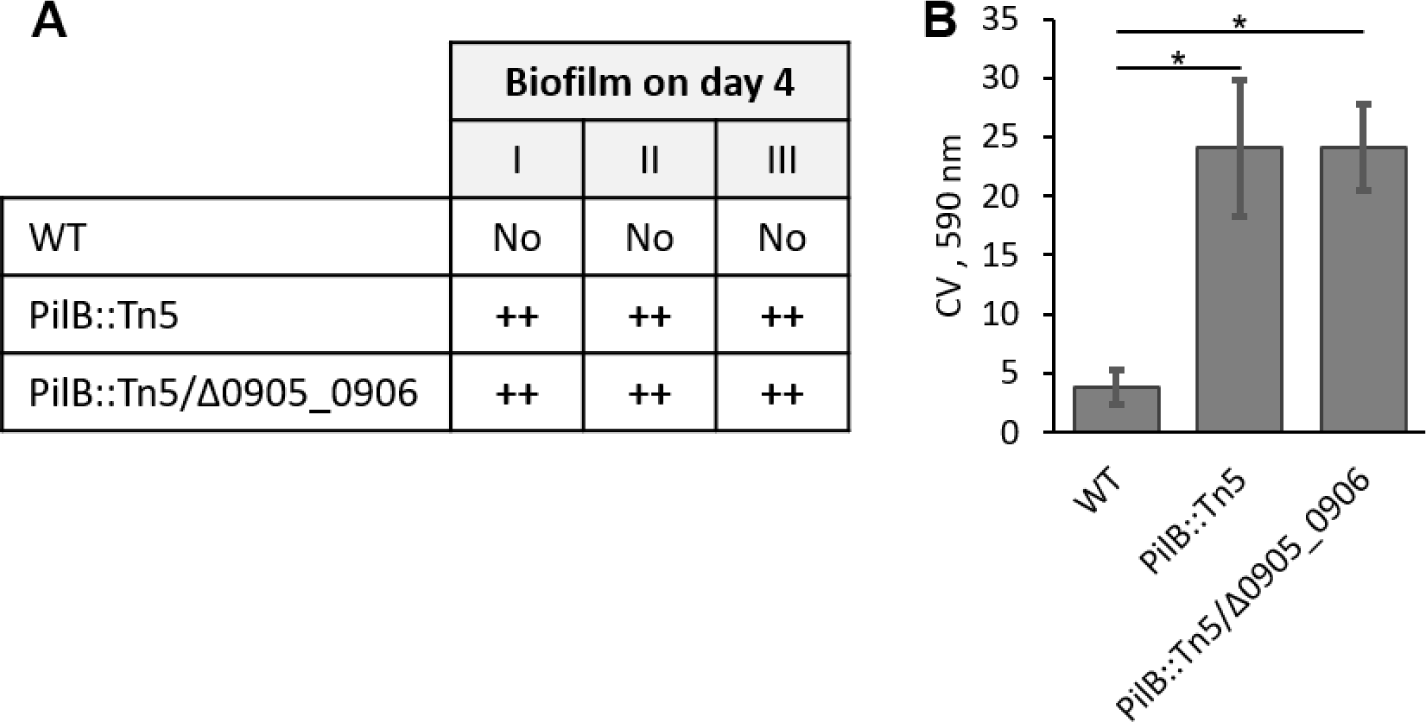
Assessment of biofilm formation under bubbling and in standing cultures in WT, PilB::Tn5 and PilB::Tn5/Δ0905_0906. **A**. Qualitative analysis of biofilm formation under bubbling (three independent experiments). **B.** Biofilm quantification by crystal violet staining in standing cultures. Asterisk denotes statistical significance (t-test, two tails, two-sample assuming unequal variances; p-value < E^-5^).

**Table S1:**
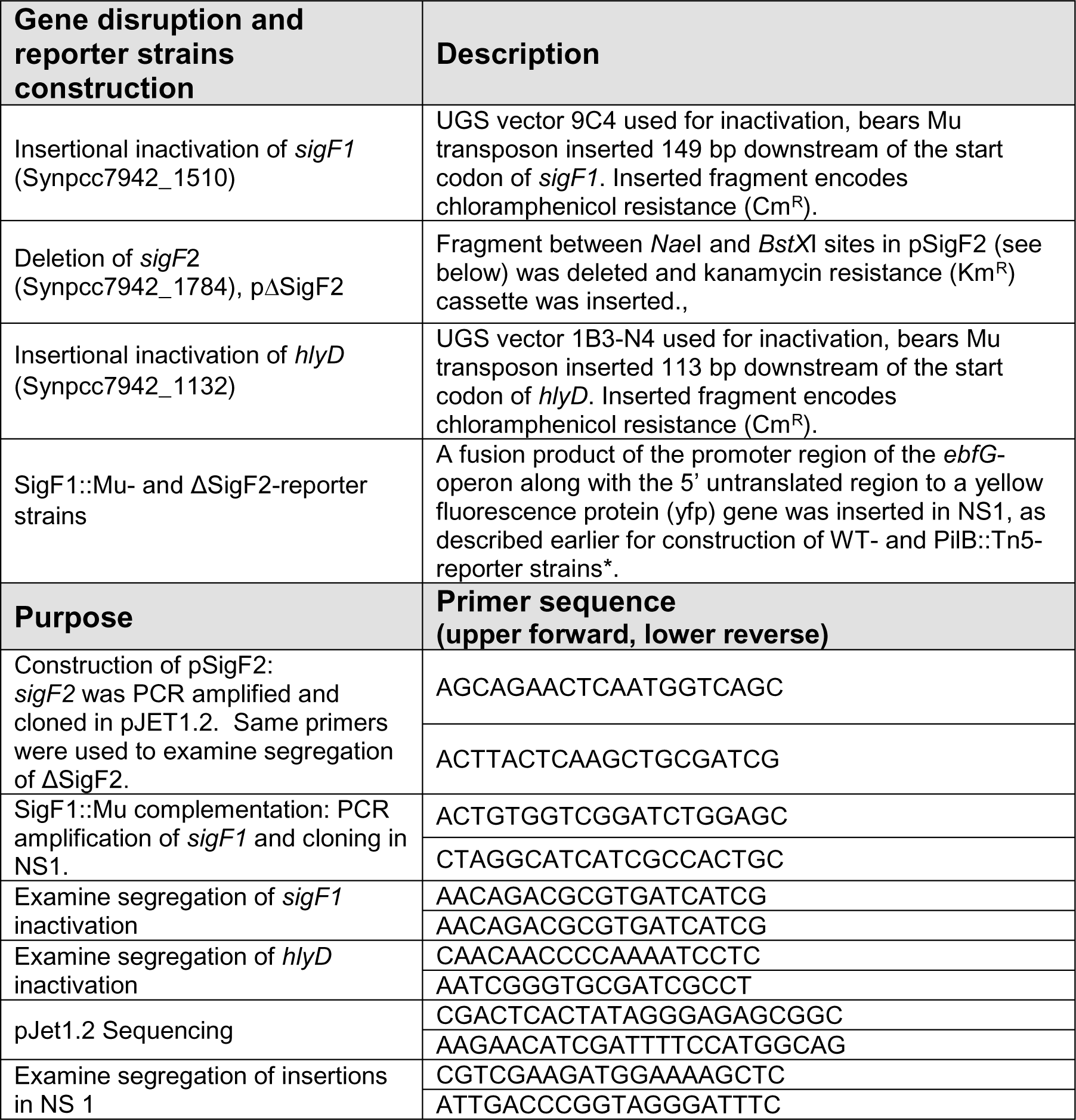
Summary of cloning information and molecular analysis. **\*** Frenkel, A., et al., *Cell specialization in cyanobacterial biofilm development revealed by expression of a cell-surface and extracellular matrix protein.* NPJ *Biofilms Microbiomes*, 2023. **9**(1): p. 10 doi.org/10.1038/s41522-023-00376-6

**Table S2:** Cofitness data for *sigF2* mutants.

| Gene | Name | Description/Annotation | Cofitness |
| --- | --- | --- | --- |
| 0001 | *dnaN* | DNA polymerase III subunit beta | 0.74 |
| 1820 |  | hypothetical protein | 0.74 |
| 0234 |  | hypothetical protein | 0.73 |
| 1668 | *kdpA* | potassium-transporting ATPase subunit A | 0.72 |
| 0619 |  | hypothetical protein | 0.71 |
| 0938 |  | transcriptional regulator, ArsR family | 0.71 |
| 2543 |  | phage SPO1 DNA polymerase-related protein | 0.71 |
| 1587 |  | integral membrane protein-like | 0.71 |
| 1607 | *somA2* | probable porin; major outer membrane protein | 0.69 |
| 0994 |  | hypothetical protein | 0.69 |
| 0500 |  | hypothetical protein | 0.68 |
| 1113 |  | hypothetical protein | 0.68 |
| 2470 |  | hypothetical protein | 0.67 |
| 1914 |  | hypothetical protein | 0.67 |
| 1942 | *prxQ-A3* | bacterioferritin comigratory protein-like | 0.67 |
| 1183 |  | hypothetical protein | 0.66 |
| 2550 |  | hypothetical protein | 0.66 |
| 1867 |  | Deoxyribodipyrimidine photo-lyase family protein | 0.65 |
| 0089 | *clcD* | Carboxymethylenebutenolidase | 0.64 |
| 1678 |  | hypothetical protein | 0.64 |

**Supplemental data file 1** – Transcriptome analysis

**Supplemental data file 2** – Exoproteome analysis

